# Structural dynamics determine voltage and pH gating in human voltage-gated proton channel

**DOI:** 10.1101/2021.08.25.457625

**Authors:** Shuo Han, Sophia Peng, Joshua Vance, Kimberly Tran, Nhu Do, Decker Gates, Nauy Bui, Zhenhua Gui, Shizhen Wang

**Author notes:** Correspondence should be addressed to SW.

## Abstract

Voltage-gated ion channels are key players of electrical signaling in cells. As a unique subfamily, voltage-gated proton (Hv) channels are standalone voltage sensors without separate ion conductive pores. They are gated by both voltage and transmembrane proton gradient (i.e ΔpH), serving as acid extruders in most cells. Amongst their many functions, Hv channels are known to regulate the intracellular pH of human spermatozoa and compensate for the charge and pH imbalances caused by NADPH oxidases in phagocytes. Like the canonical voltage sensors, the Hv channel is a bundle of 4 helices (named S1 through S4), with the S4 segment carrying 3 positively charged Arg residues. Extensive structural and electrophysiological studies on voltage-gated ion channels generally agree on an outwards movement of the S4 segment upon activating voltage, but the real time conformational transitions are still unattainable. With purified human voltage-gated proton (hHv1) channel reconstituted in liposomes, we have examined its conformational dynamics at different voltage and pHs using the single molecule fluorescence resonance energy transfer (smFRET). Here we provided the first glimpse of real time conformational trajectories of the hHv1 voltage sensor and showed that both voltage and pH gradient shift the conformational dynamics of the S4 segment to control channel gating. Our results suggested the biological gating is determined by the conformational distributions of the hHv1 voltage sensor, rather than the conformational transitions between the presumptive ‘resting’ and ‘activated’ conformations. We further identified H140 as the key residue sensing extracellular pH and showed that both the intracellular and extracellular pH sensors act on the voltage sensing S4 segment to enrich the resting conformations. Taken together, we proposed a model that explains the mechanisms underlying voltage and pH gating in Hv channels, which may also serve as a general framework to understand the voltage sensing and gating in other voltage-gated ion channels.

## Introduction

Voltage-gated cation channels mediate the electrical excitability of many cells, such as cardiomyocytes, muscular and nervous cells. As a unique subfamily, voltage-gated proton (Hvs) are solely composed of voltage sensors without the separate pore-forming domains typically found in the canonical voltage-gated cation channels (*1, 2*). Hv channel voltage sensor is a bundle of 4 helices (named S1 through S4) and ‘hourglass’ shaped, similar to these of the voltage-gated cation channels (*3, 4*). The S4 segment of the voltage-gated cation channels is highly conserved and contains multiple positively charged residues to sense membrane voltage(*5*). High resolution structures of the voltage sensors from different organisms, including Hv channels, largely agree on a model where the charge carrying S4 segment is driven outwards by depolarization voltage – a movement that is then transduced to gate the channel pore(*6*). Despite many years of research, the conformational trajectories of the voltage sensor, as well as the underlying mechanisms by which voltage and pH drive these conformational transitions in Hv channels, remain unclear, sometimes even controversial. For example, accessibility assays showed that only the middle Arg in the S4 segment of the Hv channel switches its accessibility upon voltage activation(*7*), supported by the crystal structure of a voltage sensitive phosphatase and the electron paramagnetic resonance data of the human Hv channel (hHv1)(*8, 9*). However, the gating currents of the Hv channels and many other voltage-gated cation channels suggested approximately 3 Arg residues move across the membrane upon gating transition(*10, 11*). Moreover, a unique function-defining feature of the Hv channels is that their voltage dependences are shifted by proton concentrations at the intra- and extracellular sides (i.e transmembrane pH gradient) (*12*). As a result, Hv channels are only activated when the proton electrochemical gradient is directed outwards(*13*). However, it is unclear how pH modulates the voltage sensitivity of the Hv channels. Titratable residues were proposed to be involved in the pH regulation of Hv channels and H168 was identified as a major intracellular pH sensor in the hHv1 channel(*14, 15*). In addition, the most recent studies indicated that both pH gradient and voltage alter the dynamics of the Hv channel gating currents that reflect the movements of the S4 segment (*16, 17*).

To understand the structural basis underlying voltage and pH gating in the hHv1 channel, we utilized the single molecule fluorescence resonance energy transfer (smFRET) approach that directly visualized the real time conformational transitions and dynamics of hHv1 channels in lipid environments. We performed the smFRET measurements under the voltages and pH gradients that the hHv1 gating status has been unambiguously defined. Our data clearly show that both voltage and pH gradient modify the conformational dynamics of the S4 segment in the hHv1 channel, rather than directly driving the conformational transitions between the presumptive ‘resting’ and ‘activated’ states. Our studies provide a fundamental mechanistic rationale for the pH and voltage gating in Hv channels, and probably for other voltage-gated cation channels as well.

## Results

### 1. Conformational dynamics of the hHv1 channels driven by voltages

The codon-optimized hHv1 WT channels with N-terminal 6x histidine tag were expressed and purified from *E. coli* host cells, which exhibits a single, symmetric peak in size exclusion chromatography profile (Fig S1a). The hHv1 WT protein was reconstituted into liposomes (POPE/POPG=3/1, w/w, protein/lipid = 1/200, w/w) and its channel function was examined by liposome flux assay (Fig S1b). Our results indicated that the purified hHv1 WT protein retains its native proton channel function (Fig S1c).

To implement smFRET studies on the hHv1 channel, we mutated the two intrinsic Cys107 and Cys249 residues into Ser, then introduced cysteine mutations, one pair at a time, at different sites of the hHv1 channel. Based on the available Hv channel structures (*3, 4, 18*), FRET measurements between these labeling sites can report the relative movements among the 4 transmembrane segments (Fig 1a). The hHv1 channel mutants were expressed, purified, and then labeled with Cy3/Cy5 FRET fluorophore pair with long lifetimes(*19*). The fluorophore-labeled hHv1 channel proteins were reconstituted into liposomes (POPE/POPG=3/1, w/w) with an extremely low protein lipid ratio of 1:4000 (w/w), so that most liposomes are either empty or only contain one hHv1 channel. The orientation of the hHv1 channel in proteoliposomes was controlled by incubating the proteoliposomes on the PEG passivated coverslip surface coating with anti-Histag antibodies, so only retaining these with N-terminal His-tag of the hHv1 facing outside. Thus, the defined electrical potential can be generated with K^+^ gradients and K^+^ ionophore valinomycin and applied to the hHv1 channel in liposomes (Fig 1a). The smFRET imaging was performed on a customized TIRF microscope with a dual-view beam splitter and EMCCD camera as described previously(*20, 21*), and the image data were processed by the SPARTAN software (*22*). The liposome voltages can be calculated with the Nernst equation and switched by changing the extraliposomal K^+^ concentrations (Fig 1a). Our initial smFRET measurements did not detect any significant voltage-dependent FRET changes at the voltage sensing S4 segment (Fig S2a). We then realized that the proton uptake through the hHv1 channel will decrease the extracellular pH of the hHv1 channel (i.e the intraliposomal pH) and deactivate it in less than 1 min (Fig S2b), therefore the activated conformations are too short to be captured. We then introduced the N214R mutation, which was showed to abolish the H^+^ conductance without altering the voltage dependence of the hHv1 channel (*10, 11*). Our liposome flux assay also confirmed that the hHv1 N214R mutant protein exhibits no visible proton uptake (Fig S2b). We further confirmed that, most fluorophore labeled hHv1 mutant proteins for smFRET studies remain functional, in the absence of the N214R mutation (Fig S1c). On the N214R background, substantial voltage-dependent FRET changes were detected at both the K169C-Q194C and K125C-S224C labeling sites, which are predicted to report the relative movements of the S4 segments, but with reversed FRET changes (Fig 1a). SmFRET traces from the K169-Q194 labeling sites, exhibit large, spontaneous transitions among three major FRET states with peaks at 0.24, 0.52 and 0.78, respectively, at both resting (−85 mV) and activating voltages (Fig 1b, Fig S3a). For all these traces, the donor and acceptor fluorescence intensities are strongly anticorrelated, indicated the FRET changes truly result from structural changes rather than fluorophore photophysics (Fig 1b, Fig S3a). Histograms and contour maps from several hundreds of hHv1 channels indicated that weak (0 mV) and strong activating (120 mV) voltages significantly enrich the low FRET-0.24 populations, but diminish the medium and high FRET populations (Fig 1c, f). Shifts of the FRET population indicated that activating voltage enhances the stabilities of the S4 segment conformations facing outwards. These results were further validated by these from the K125C-S224C sites (Fig 1a). Similarly, large spontaneous FRET transitions were also observed under all voltages with the activating voltages (0 and 120 mV), in sharp contrast, promoting the high FRET populations (Fig 1d, Fig S3b). The voltage-dependent shifts of three major FRET populations with peaks at 0.27, 0.6 and 0.9 are well demonstrated by the histograms and contour maps from >600 molecules (Fig 2e, g). Together, these smFRET data consistently indicate that the voltage sensing S4 segment of the hHv1 voltage sensor undergoes spontaneous transitions. The activating voltage only enriches, but does not directly drive the S4 segment toward the outward positions, i.e the conformations that are often defined as the activated state in X-ray or cryo-EM structures.

**Fig 1.**
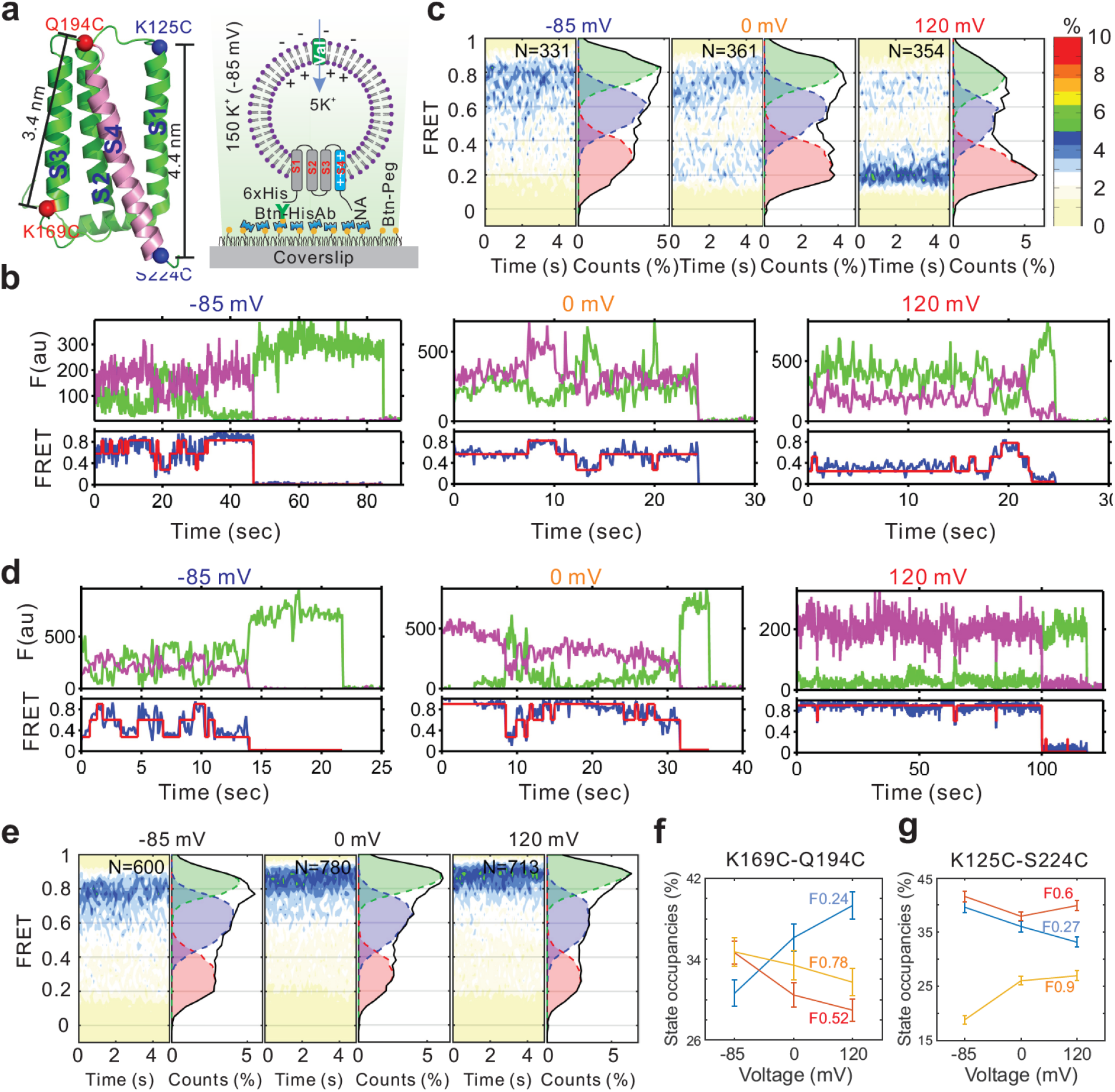
The voltage-dependent conformation dynamics of the hHv1 channel S4 segment revealed by smFRET. a. Left panel, the cartoon of the hHv1 NMR structure (5oqk, left panel) with the alpha carbons of the K125C-S224C (blue) and K169C-Q194C (red) labeling sites highlighted by spheres; Right panel, the sample immobilization configuration for smFRET imaging. The PEG passivated surface with 2% PEG-biotin binds the biotinylated anti-Histag antibodies (Btn-HisAb) via neutravidin (NA), then the hHv1 liposomes with N-terminal 6*Histag at outside were selectively retained for smFRET imaging under different voltages. b, d. Representative smFRET traces between the K169C-Q194C and K125C-224C labeling sites under -85 (resting), 0 (weak) and 120 mV (strong activating) voltages. The green and pink lines are donor and acceptor fluorescence intensities, the blue and red lines are the real and idealized FRET. c, e. FRET histograms and contour maps of the smFRET data obtained from the K169C-Q194C, K125C-224C labeling sites under different voltages. f, g. State occupancies calculated from the smFRET data between the K169C-Q194C and K125C-224C labeling sites under different voltages.

**Fig 2.**
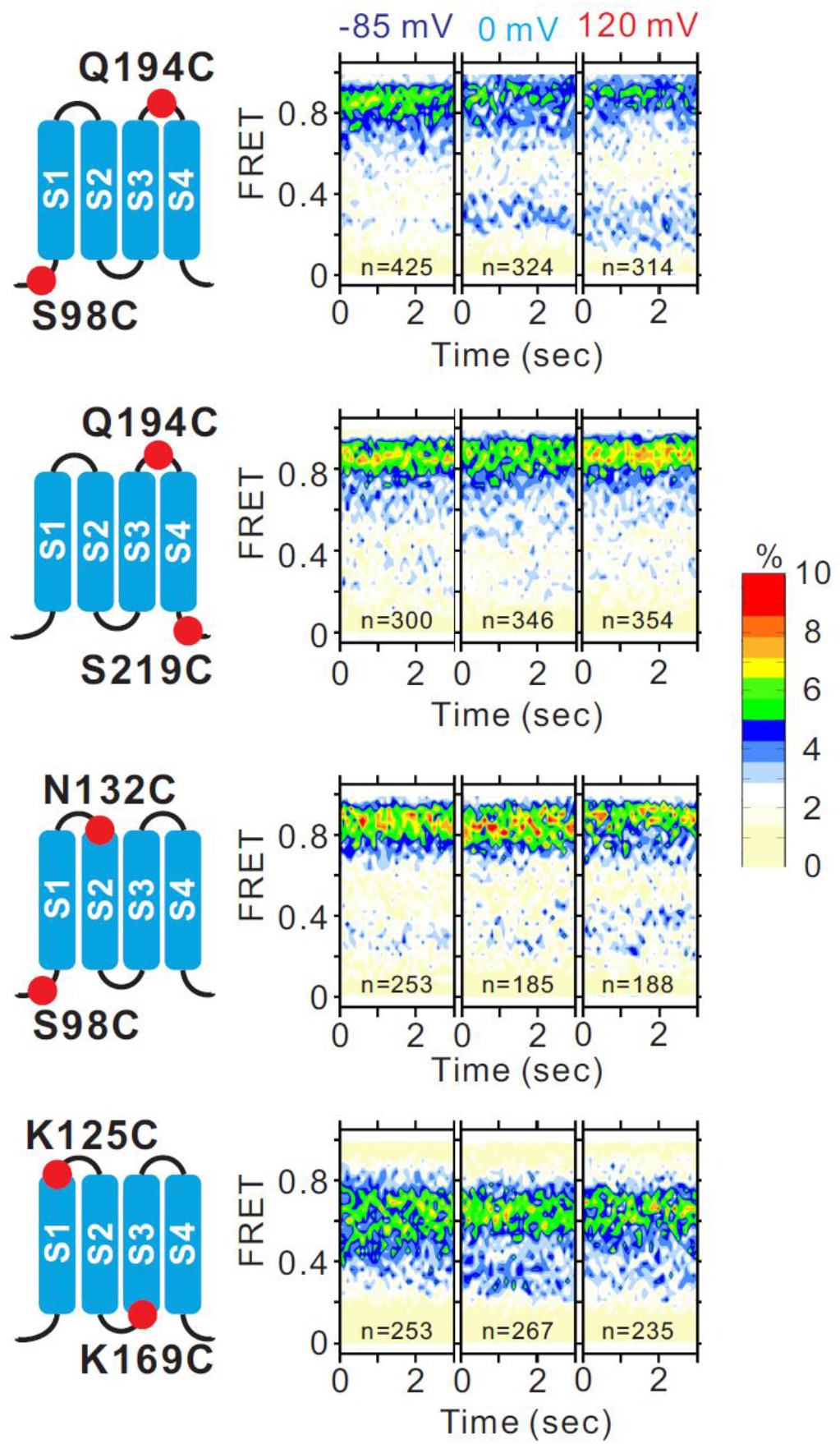
Global conformational distributions of the hHv1 voltage sensor at different voltages. SmFRET contour maps were calculated from the smFRET data of the first 5 secs between the a. S98C-Q194C, b. Q194C-S219C, c. S98C-N132C, d. K125C-K169C labeling sites, the bin size is 0.03.

### 2. Global conformational changes of the hHv1 channels at resting and activated states

To map the global conformational changes of the hHv1 channel voltage sensor, smFRET measurements were performed systematically at multiple labeling sites, one pair at a time, at different voltages. SmFRET contour maps showed that the S98C-Q194C sites (S1 vs S4) have similar voltage-dependent changes as that of K169C-Q194C sites, while the Q194C-S219C sites (S4 itself), S98C-N132C (S1 vs S2) and K125C-K169C sites (S1 vs S3) did not exhibit any voltage-dependent shifts (Fig 2). These data indicate that activating voltages only act on the voltage sensing S4 segment to promote the outwards conformations, while the S1, S2 and S3 segments do not exhibit detectable movements (Fig 2, Fig S3c).

Taken together, our key finding is that the S4 segment appears to be highly dynamic and can reach all the three FRET states, even at the resting voltage of -85 mV (Fig 1b, d). Our smFRET data suggest that voltage determines the conformational dynamics of the S4 segment in a probabilist, rather than deterministic manner. In other words, in terms of the voltage-sensing S4 segment, the opening or closing states are the same groups of conformations, and it is the differences in the conformational distributions that determine the biological gating status. The small shifts in FRET distributions may reflect the small open probability changes of the hHv1 channels at the voltages and pHs tested in the present work, for the maximum open probability of the hHv1 channels in human eosinophils is ∼ only 0.6 (*23*).

### 3. pH gates hHv1 by shifting the conformation dynamics of the S4 segment

To understand how pH shifts the voltage sensitivity of the hHv1 channel, smFRET measurements were performed on the K125C-S224C sites at different pH (Fig 3a). At 0 mV, lower the intracellular pH (i.e the extraliposomal pH) to 6.5 remarkably enhances the high FRET populations of the S4 segment, consistent with its effect to shift the voltage dependence more negatively(*12*) (Fig 3a, left). In comparison, changing both the intra- and extracellular pHs symmetrically does not impact the FRET distributions of the S4 segment (Fig 3a, middle), matching with the electrophysiological results that it is the transmembrane pH gradient that regulates the hHv1 channel gating(*12*). To further test whether the pH effects on the S4 segment are specific and related to channel gating, we introduced H168Q mutation in the hHv1 channel that abolishes its intracellular pH dependence (*15*). The smFRET data from the same K125C-S224C sites on the H168Q mutation background indicate that the high FRET populations are significantly increased, and no longer shifted by the intracellular pHs (Fig 3a, right). Our data conclusively indicate that pH gating of the hHv1 channels originates from the pH-dependent conformational changes of the voltage sensing S4 segment.

**Fig 3.**
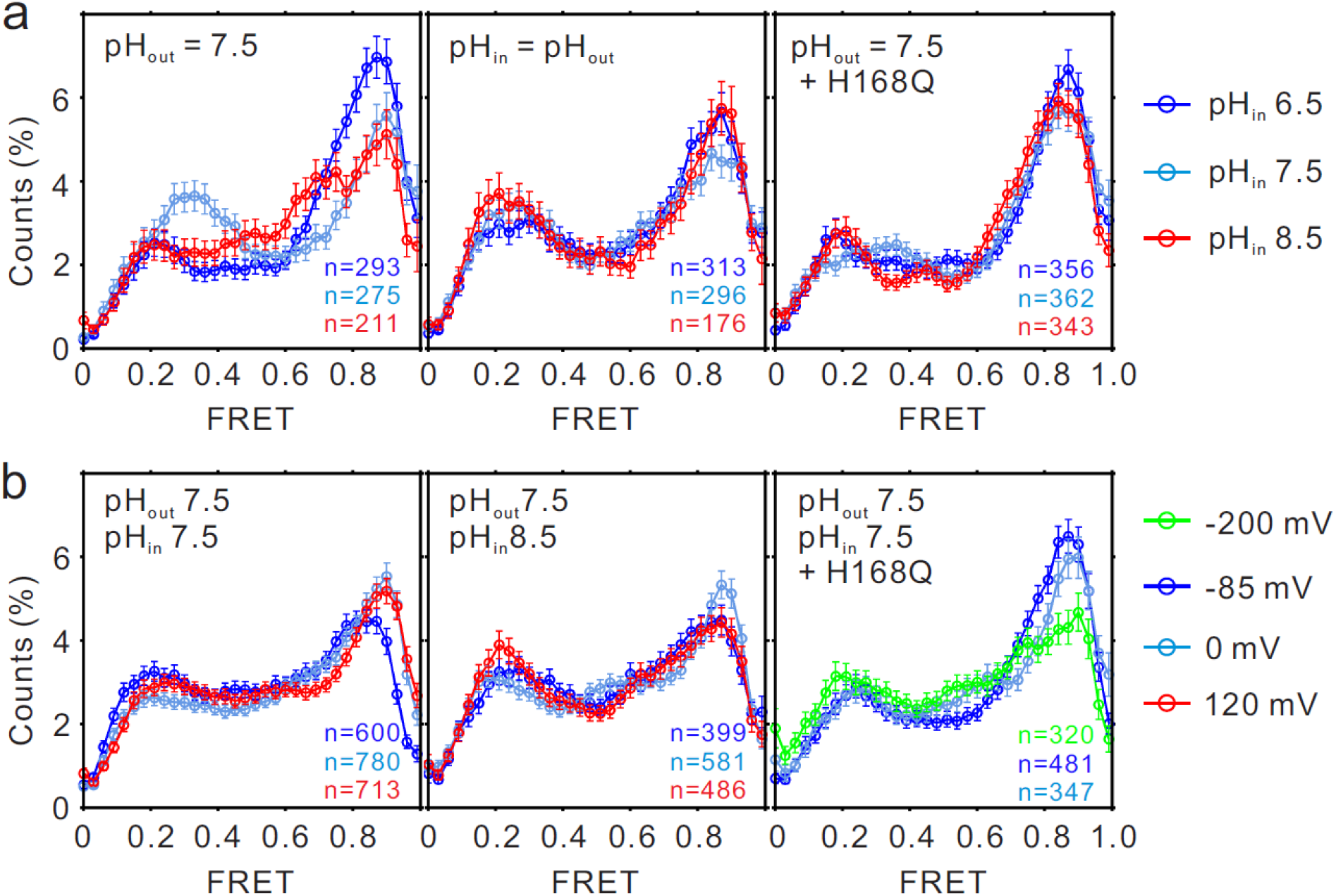
The structural basis underlies pH regulating the voltage dependence of the hHv1 channels. a. Histograms of the smFRET data from the K125C-S224C sites of the hHv1 channels in liposomes under 0 mV with asymmetrical (left and right panels), symmetrical (middle) pH conditions on the WT (left, middle panels) or H168Q mutation backgrounds. b. Histograms of the smFRET data from the K125C-S224C labeling sites of the hHv1 channels on the WT (left, middle panels) and H168Q mutation backgrounds with or without pH gradient at different voltages.

To further elucidate the pH voltage interplay in determining hHv1 channel gating, we elevate the intracellular pH of the hHv1 channel to 8.5, which shifts the voltage dependence positively. Consistently, the enrichments of the high FRET populations by activating voltages were undetectable on the K125C-SS24C labeling sites (Fig 3b, right and middle). H168Q mutant under 0 mV demonstrates very high FRET populations (Fig 3a, right), predictin that it may require a more negative voltage to be stabilized at resting conformations. Indeed, we found that the low FRET populations correlating with channel closure are only observed on K125C-S224C labeling sites at more negative -200 mV on the H168Q, not at -85 mV on the WT background (Fig 3b, right). Our results suggest that H168 interacts with the S4 segment to stabilize it at inward conformations that endorse channel deactivation.

### 4. Identification of the extracellular pH sensor in the hHv1 channel

Previously we introduced the N214R mutation to prevent the deactivation of the hHv1 channel by extracellular pH (Fig 4a), which permitted us to capture the activated conformations (Fig 1c, e). In the absence of the N214R mutation, the FRET distributions of the K169C-Q194 sites at all voltages are dominated by the high FRET populations, i.e the conformational distributions appearing at resting voltage (Fig 4b). However, if the extracellular pH sensor/s can be removed, the ‘activated’ FRET distributions can be captured even the hHv1 channel is still functional to decrease extracellular pH (i.e intraliposomal pH, Fig 4a). We looked into the hHv1 channel structures and found H140 is very close to the S4 segment, which may serve as the extracellular pH sensor/s. The H140A mutation was introduced to the K169C-Q194C mutant and liposome flux assay indicates that the resulting mutant remains functional to mediate proton uptake into liposomes (Fig 4c). On the H140A mutation background, the FRET distributions at the K169C-Q194C labeling sites do shift by voltage (Fig 4d), the same pattern as that on the N214R background (Fig 1c). But, we do notice that the low FRET populations at 120 mV are slightly less enriched than that on the N214R background, suggesting that H140 perhaps is not the only extracellular pH sensor of the hHv1 channel. Similar results were also observed on the K125CS224C labeling sites (Fig S4a, b), further confirmed that H140 is the key residue in the hHv1 channel that senses extracellular pH changes.

**Fig 4.**
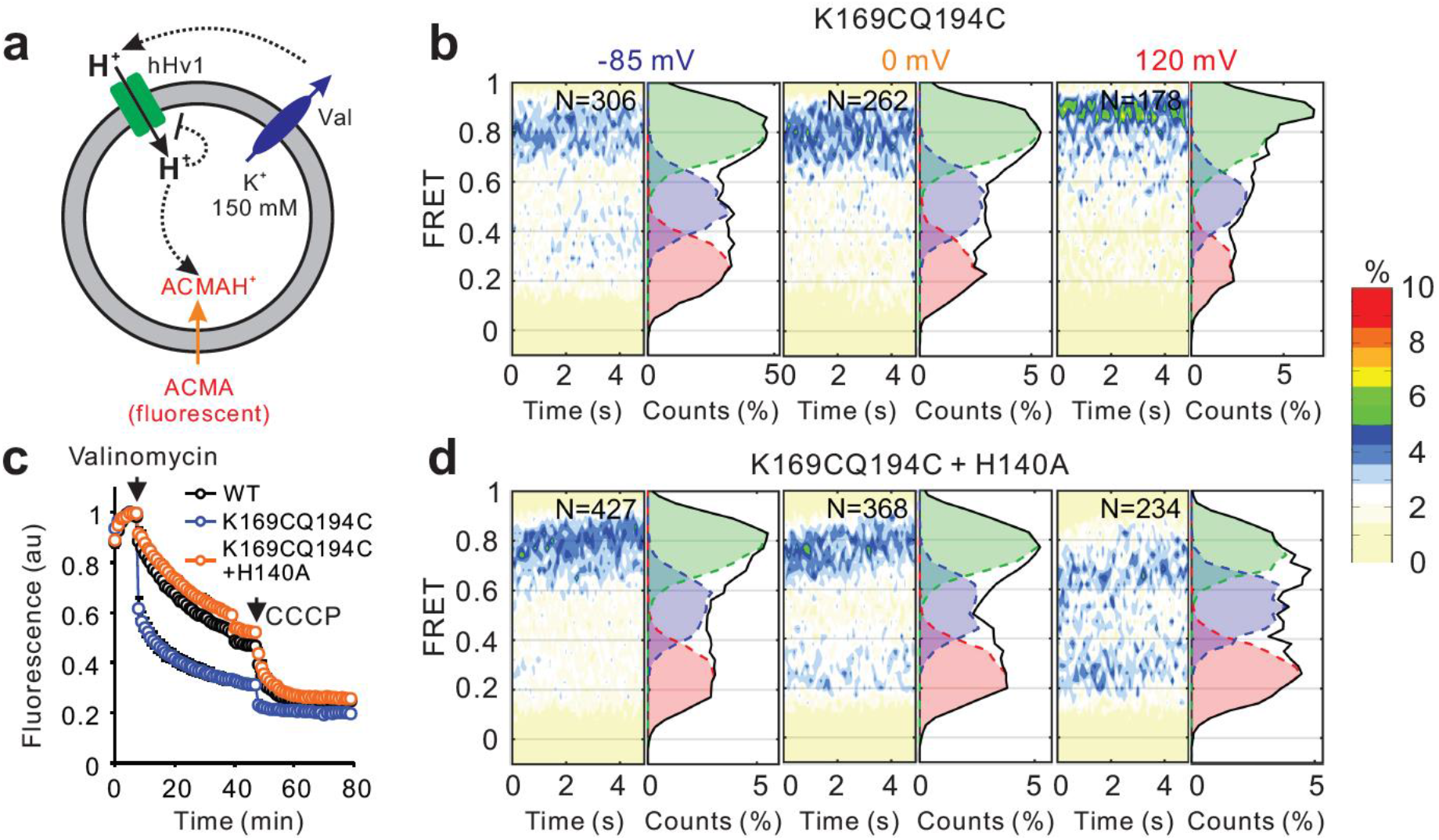
H140 is the extracellular sensor in the hHv1 channel. a. Proton uptake into liposomes through the functional hHv1 channel deactivates it by lowering the extracellular pH, indicating by quenching of ACMA fluorescence. b, d The histograms and contour maps of the K169CQ194C mutant without (c) or with (d) H140A mutation that removes proton inhibition of the hHv1 channel at the extracellular side. c. Liposome flux data showed that the WT, K169CQ194C and K169CQ194C-H140A mutants are all functional to mediate proton uptake. The addition of valinomycin at ∼7 min triggers K^+^ outflow following chemical gradient to generate electrical potential, which drives proton uptake into liposomes and quenching of the ACMA fluorescence.

In summary, our results suggest that both pH and voltage act on the S4 segment of the hHv1 voltage sensor to modulate channel gating, which is consistent with the most recent gating current recordings(*17*). Based on our smFRET data, we proposed a gating model, as shown by Fig 5, which may underline the pH and voltage gating in the hHv1 channels. The intrinsical threshold activating voltage of the S4 segment in the hHv1 voltage sensor probably is much more negative, like these from canonical voltage-gated ion channels. But, the pH sensors at the intra (H168) and the extracellular (H140) sides physically interact with the S4 segment to stabilize it at inwards conformations that endorse channel deactivation. The molecular couplings between the S4 segment and pH sensors are dynamic and dependent upon the protonation state of the H168 and H140 residues. Meanwhile, the S4 segment undergoes spontaneous upwards and downwards transitions, which are dependent on the membrane voltage. Although the extra and intracellular pHs do not directly act on the voltage sensing S4 segment, they modify the dynamics of molecular couplings between the pH sensors and S4 segment. Thus the occupancies of the preactivated and activated conformations are shifted to change the voltage dependence, either negatively by destabilizing, or positively by stabilizing the molecular couplings between the pH sensors and the S4 segment in the hHv1 channel.

**Fig 5.**
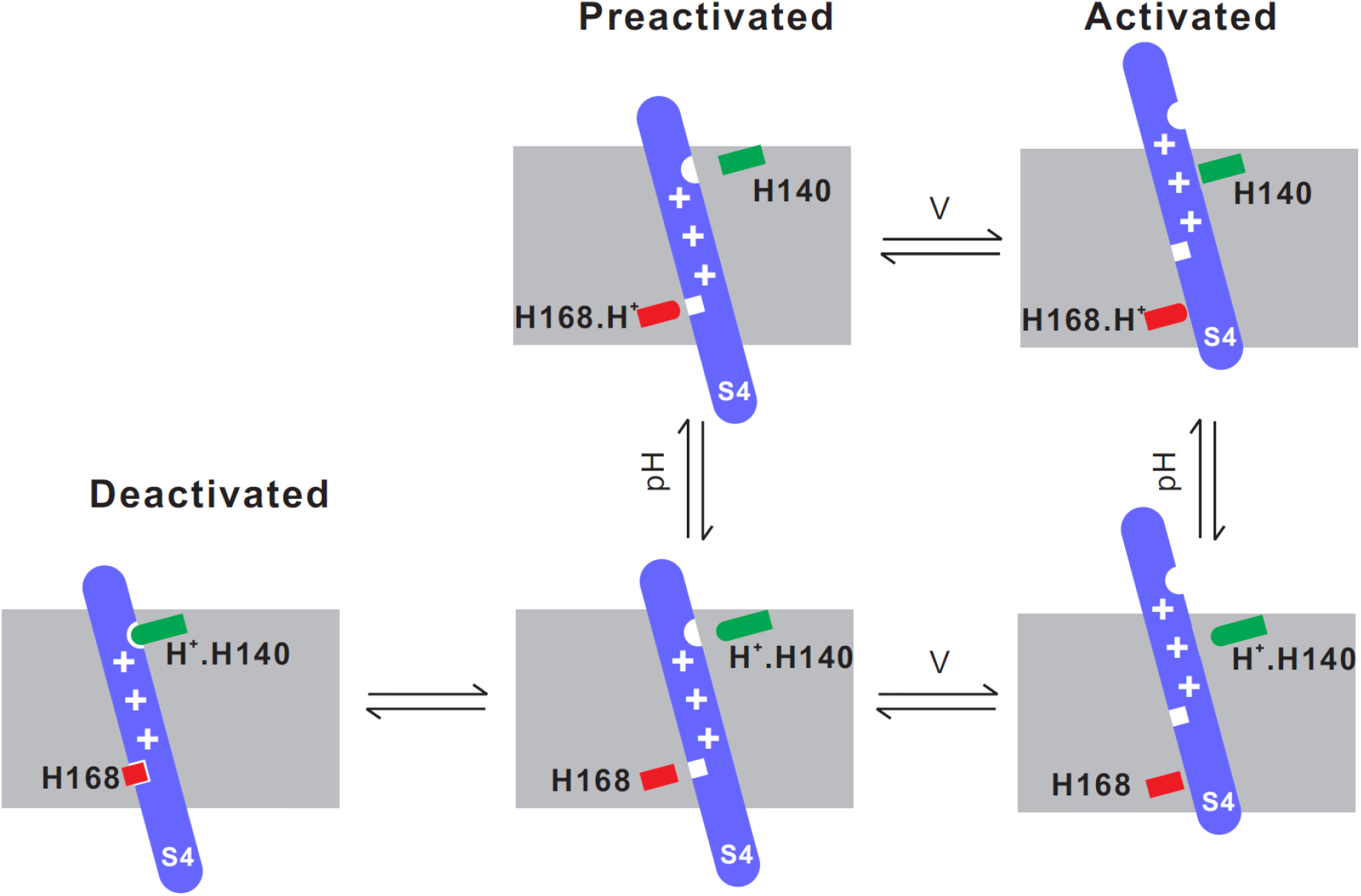
The molecular model to explain the voltage pH interplay in gating the hHv1 channels. The H168 (red) and protonated H140 (green) at the extracellular side lock the S4 segment (blue) of the hHv1 voltage sensor at the intracellular side. The interactions between the pH sensors and the S4 segment are dynamic, which are shifted by both the intra- and extracellular pHs. Decoupling of the S4 segment from pH sensors allows the spontaneous transitions between the inwards and outwards conformations, which are dependent on membrane voltage.

## Discussion

Structural alignments of the voltage sensors from different organisms or even the same channels largely agree on a model where the positively charged S4 segment is driven outwards by depolarization – a movement utilized by pore modules to control ion permeation(*6, 9, 24*). However, the details of this movement are far from clear, different movements have been described for different voltage sensors. For example, the S4 segment moves up for 3 helical turns in a voltage-gated sodium channel NavAb (*24*) while only 1 helical turn in a voltage sensitive phosphatase(*9*). Unfornaturely, most atomic structures of voltage-gated ion channels were obtained at one voltage (i.e 0 mV in solution) and their gating status can not be determined, even enter conformations that may not be physiologically relevant (*25*). Thus the conformational changes of the voltage sensors inferred from the structural alignments are yet to be functionally defined. We performed quantitative analyses on the real time conformational transitions and dynamics of the hHv1 voltage sensor in lipid environments, at voltages and pHs that the hHv1 channel functional status has been clearly defined. Our results showed voltage and pH modify the voltage sensor conformations of the hHv1 by shifting the dynamic distributions between the conformations endorsing either deactivated or activated states, in a probabilistic, rather than deterministic manner. The gating currents Hv channels suggested multiple close states (*10, 26*), consistently, the S4 segment exhibits multiple conformations under a strong resting voltage of - 85 mV in our studies. Since the fluorophores were attached to the well-exposed residues in the hHv1 channel with relatively long, flexible C5 linkers, they probably adopt random orientations that sample the cone spaces (i.e K^2^=2/3). The large, spontaneous movements of the S4 segment predicted by smFRET could be slightly over 20 angstroms, which suggests that the S4 segment probably moves at least 3 helical turns (i.e 3×5.4 Å) to reach activating states. At both strong resting and activating voltages, the S4 segment transits among the same group of conformations and it is the conformational distributions that are highly correlated with the gating voltages. Intriguingly, the spontaneous, large conformational transitions of the S4 segment in the hHv1 channel perhaps are the structural basis of the gating current noise (*27*). In conclusion, the unique voltage and pH gating in the hHv1 channel is based on the spontaneous, voltage-dependent structural transitions of the S4 segment, which facilitate its pH dependent dynamic interactions with both the intra-(H168) and extracellular (H140) pH sensors. Our gating model (Fig 5) also explains the most recent patch-clamp fluorometry data and gating current from Hv channels, which showed that the conformation transitions of the S4 segment are dependent on both intra- and extracellular pHs (*17, 28*).

## Abbreviations

POPE: 1-palmitoyl-2-oleoyl-sn-glycero-3-phosphoethanolamine
POPG: 1-palmitoyl-2-oleoyl-sn-glycero-3-phospho-(1’-rac-glycerol)
TCEP: tris(2-carboxyethyl)phosphine
ACMA: 9-Amino-6-Chloro-2-Methoxyacridine
CCCP: carbonyl cyanide m-chlorophenyl hydrazine
2-GBI: 2-guanidinobenzimidazole
NADPH: reduced nicotinamide adenine dinucleotide phosphate
NMDG: N-Methyl-D-glucamine
ROS: reactive oxygen species
PCD: protocatechuate 3,4-Dioxygenase
PCA: protocatechuic acid
COT: cyclooctatetraene
Hepes: 4-(2-hydroxyethyl)-1-piperazineethanesulfonic acid
Tris: 2-Amino-2-(hydroxymethyl)propane-1,3-diol
EMCCD: electron multiplying charge coupled device

## Author contributions

SW conceived the studies, SW and SH designed the studies. SH, SP and JV performed the studies with help from KT, ND, DG, NB and ZG; SW and SH analyzed the data, SW prepared the manuscript with editing input from other authors.

## Acknowledgments

This work was funded by NIH grant 1R15GM137215-01 (SW) and the startup fund of UMKC. We would like to thank Dr. Lejla Zubcevic at the Kansas University Medical Center and Dr. Xiaolan Yao at the University of Missouri-Kansas City for their help in reviewing and revising the manuscript.

## METHODS

### 1. Protein expression, purification and fluorophore labeling

The hHv1 cDNA was codon-optimized and synthesized by Genscript Inc, then inserted into pET28a(+) vector between the NdeI and BamHI restriction sites. The resulting hHv1 protein contains a 6x Histidine tag at N-terminus, following by a thrombin protease cutting site. All mutations were introduced into hHv1 by site-direct mutagenesis (Agilent Inc.) and confirmed by DNA sequencing. The hHv1 proteins were expressed in *E. coli* BL21 (DE3) host cells and purified as that described by Li et al(*8*). In brief, the hHv1 proteins were purified by metal affinity chromatography then loaded onto a Superdex-200 size exclusion column with running buffer containing 20 mM Tris, 150 mM NaCl, 1 mM Fos-Choline 12, 1 mM TCEP, pH8.0. For smFRET imaging samples, the proteins were subjected for buffer exchange using 5 mL Hi-Trap desalting column to labeling buffer containing 20 mM Hepes, 150 NaCl, 1 mM Fos-Choline 12 (n-Dodecylphosphocholine), pH 7.0, then mixed with Cy3 and Cy5 c5 maleimide(*19*) 1:1 mixture at a protein dye molar ratio of 1:6. The labeling reactions were performed under 4 °C for 3 hours, then the free fluorophores were completely removed by performing a metal affinity chromatography and a size exclusion chromatography in a row. All the proteins were either stored at -80 °C for later use or reconstituted immediately.

### 2. hHv1 Liposome reconstitution

The lipids dissolved in chloroform containing POPE/POPG (3:1, w/w, Avanti Polar Lipids Inc.) were dried in clean glass tubes with argon gas first, and then in a vacuum desiccator for 4 hrs to evaporate organic solvent completely. The dried lipids were resuspended in reconstitution buffer containing 20 mM Hepes, 5 mM KCl, 150 mM NaCl, pH7.5 (for smFRET imaging) or 20 mM Hepes, 150 mM KCl, 0.05 mM NaCl, pH7.5 (for liposome flux assay). The liposomes were formed by sonication, then destabilized with 10 mM Fos-choline 12 (Anatrace Inc). For liposome flux assay, the purified protein was mixed with the lipid solution at a protein lipid ratio of 1:200 (w/w), and the hHv1 proteoliposomes were formed by detergent removal using Bio-beads SM2 (Bioad Inc). For smFRET imaging, the proteins were mixed with lipids at a ratio of 1:4000 (w/w) and the detergents were removed by dialyzing against reconstitution buffer containing 1 mM TCEP at the volume ratio of 1:500 for at least 3 times, each time over 12 hours.

### 3. Liposome fluorescence flux assay

The K^+^ gradient between inside and outside of the liposomes is established by diluting the liposomes in the extraliposomal buffer containing 20 mM Hepes, 150 mM NMDG, pH7.5. The liposomes were incubated with 2 µM of ACMA fluorescence probes for ∼5 min, then ACMA fluorescence was measured using a 96 well plate reader (FluoStar, Ex/Em = 390nm/460nm) for ∼5 min. After valinomycin was added at a final concentration of 0.45 µM, the fluorescence measurements were resumed with the same optical setting for ∼40 min. Proton-specific ionophore CCCP was used as the positive control, liposomes without K^+^ gradient, or with K^+^ gradients without valinomycin were used as negative controls. The fluorescence liposome flux data were processed following the method of Su et al (*29*). In brief, the hHv1 channel activities were calculated from the normalized fluorescence readings using the equation:

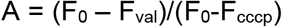

Where F_0_, F_val_ and F_cccp_ are the stead state ACMA fluorescence at initial, after adding valinomycin and CCCP, respectively. The relative activities of the hHv1 channel were normalized against the hHv1 WT proteoliposomes included in every batch of assays.

### 4. Single molecule imaging and data analysis

Flow chambers for smFRET imaging were prepared following the protocol of Joo et al (*30*). An objective-based TIRF built on a Nikon TE-2000U inverted microscope (TE-2000s) with 100x APO TIRF NA1.49 objective lens, 532 nm and 640 nm lasers, was used for single molecule imaging. Donor and acceptor emissions were separated by W-view Gemini beam splitter with chromatic aberration correction (Hamamatsu Inc.) carrying the 638 nm long-pass beam splitter, then cleaned by 585/65 nm and 700/75 nm bandwidth filters (Chroma Inc.). The images were collected by an ImagEM X2 EMCCD camera (Hamamatsu Inc.). The liposomes containing fluorophore-labeled hHv1 channels were retained on the PEGlyated surface coated with biotinylated Histag antibodies (1:200 dilution, ThermoFisher Inc.). Fluorophores were excited by 532 nm laser (∼1.0 W/cm^2^) and time-lapse movies were collected at 10 frames per second (i.e. time resolution of 100 ms). Typical recording times were ∼3 min, with half bleaching times ∼ 1min. All imaging buffers contained ∼3 mM Trolox, 5 mM PCA and 15 μg/μL of PCD to enhance the photostability of the fluorophores(*31, 32*). For symmetrical pH conditions, β-escine was used to permeabilize liposomes(*33*). At least 3 batches of independent smFRET imaging were performed for each sample/condition. For every movie, there were ∼500 molecules per field of view with ∼100 molecules containing both acceptor and donor fluorophores. The smFRET traces were extracted and analyzed with the SPARTAN software(*22*) and then manually picked following the criteria described in the previous studies(*34, 35*). The bin size of all histograms and contour maps was 0.03 and FRET contour plots were generated from the smFRET data of the first 5s. For kinetic analysis, a three-state FRET model was used and the smFRET traces were idealized with the SPARTAN software first using the Baum-Welch, then the Maximum Point Likelihood algorithms. The same peak centers were applied for analyzing smFRET data from the same labeling sites K125C-S224C or K169C-Q194C.

## Extended Figures

**Fig S1.**
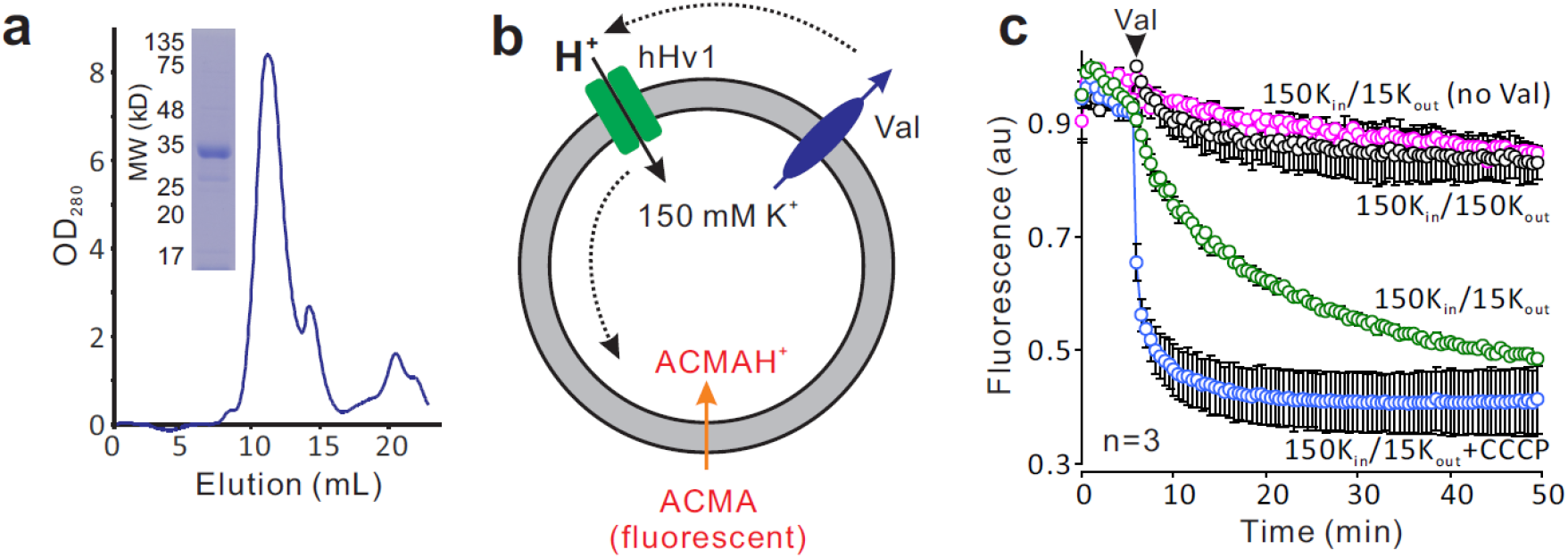
The purified hHv1 protein functions as a proton channel. a. SDS-PAGE and size exclusion chromatography profiles of the hHv1 channel WT protein expressed and purified from *E. coli* host cells. b. The liposome fluorescence flux assay to determine channel activities of the hHv1 channel. The Nernst potential is generated and can be calculated by the K^+^ gradient across the liposomes in the presence of K^+^ ionophore valinomycin (Val). Proton uptake through the hHv1 channel into liposomes quenches the ACMA fluorescences. c. Liposome flux assay of the proton uptake mediating by hHv1 channel reconstituted into liposomes (POPE/POPG = 3/1, protein/lipid = 1/200, w/w), n=3.

**Fig S2.**
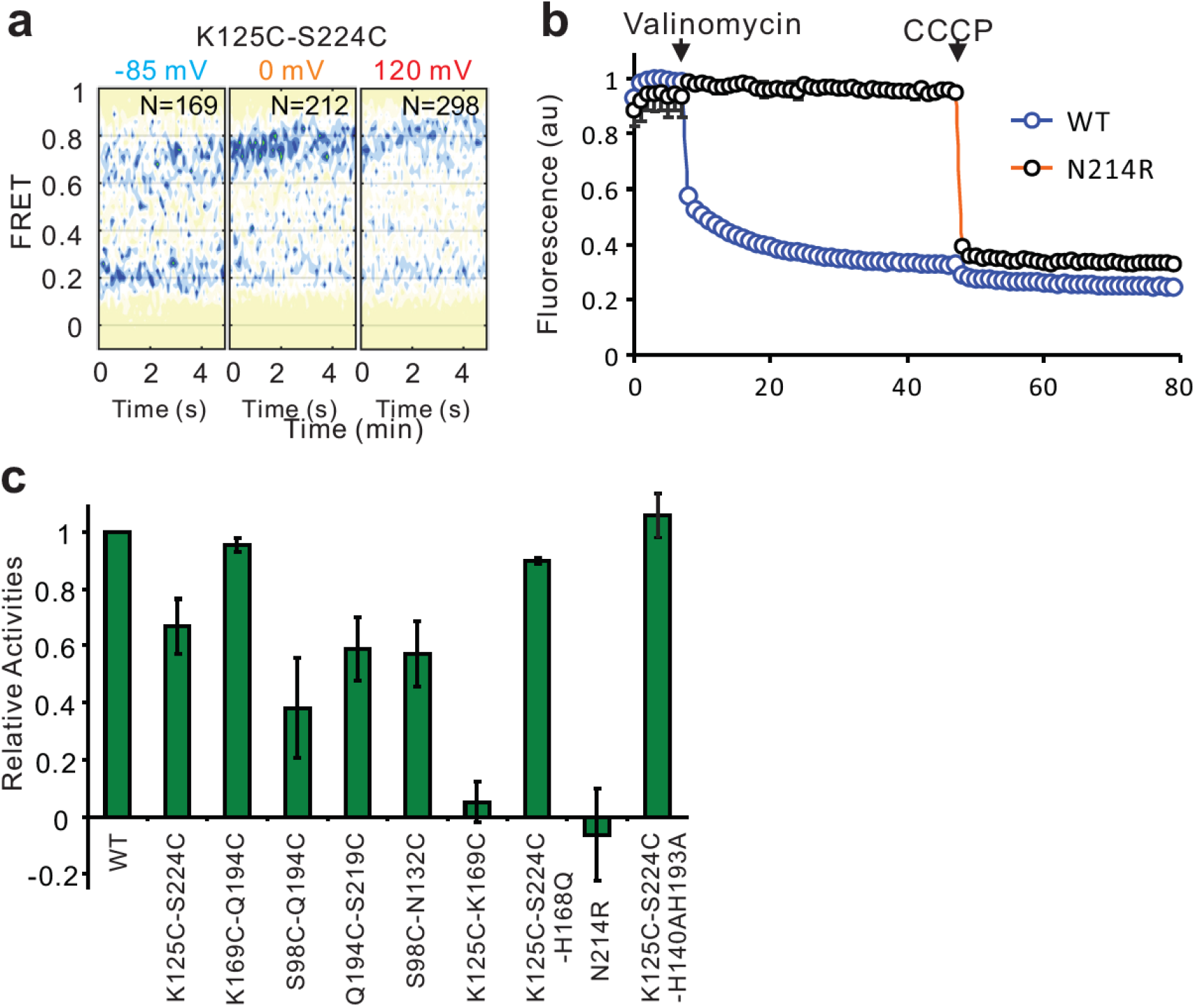
N214R mutation abolishes hHv1 channel pH inhibition by preventing proton uptake into the liposomes. a. Contour maps and histograms of the smFRET data between the hHv1 K125C-S224C sites on the WT background at different voltages. b. The liposome flux assay of hHv1 WT and hHv1-N214R mutant proteins. The arrows marked the time points when the valinomycin (0.45 µM) and CCCP (1 µM) were added. c. Relative channel activities of the fluorophore-labeled hHv1 mutant proteins subjected to smFRET studies, without the N214R mutation, determined by liposome flux assay. For all samples, n=3.

**Fig S3.**
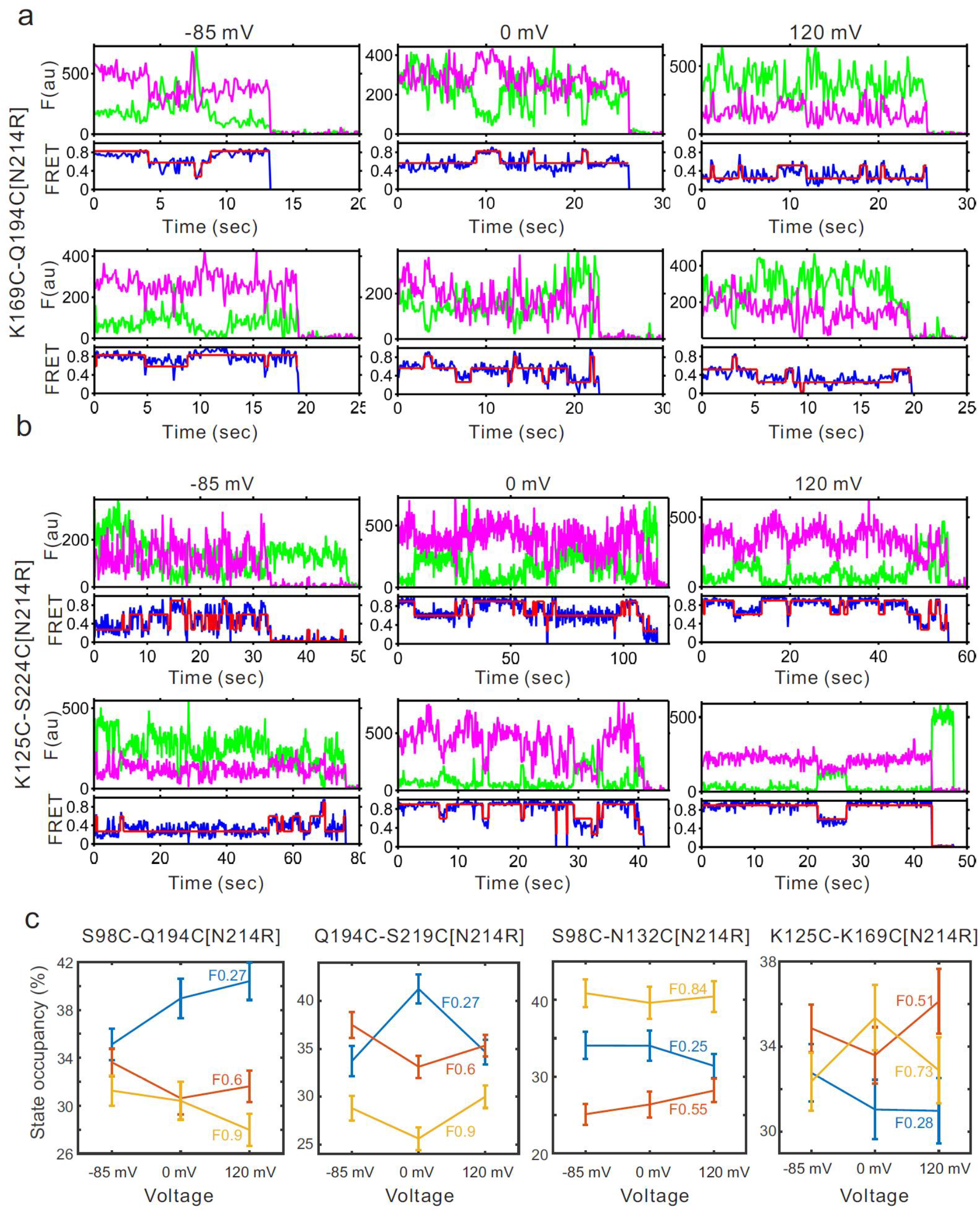
Conformational changes of the hHv1 channels revealed by smFRET. a&b, representative smFRET traces of the K169C-Q194C (a) and K125C-S224C (b) labeling sites on the N214R mutation background at different voltages; c. FRET state occupancies that were calculated from the smFRET data of the S98C-Q194C, Q194C-S219C, S98C-N132C and K125C-K169C labeling sites.

**Fig S4.**
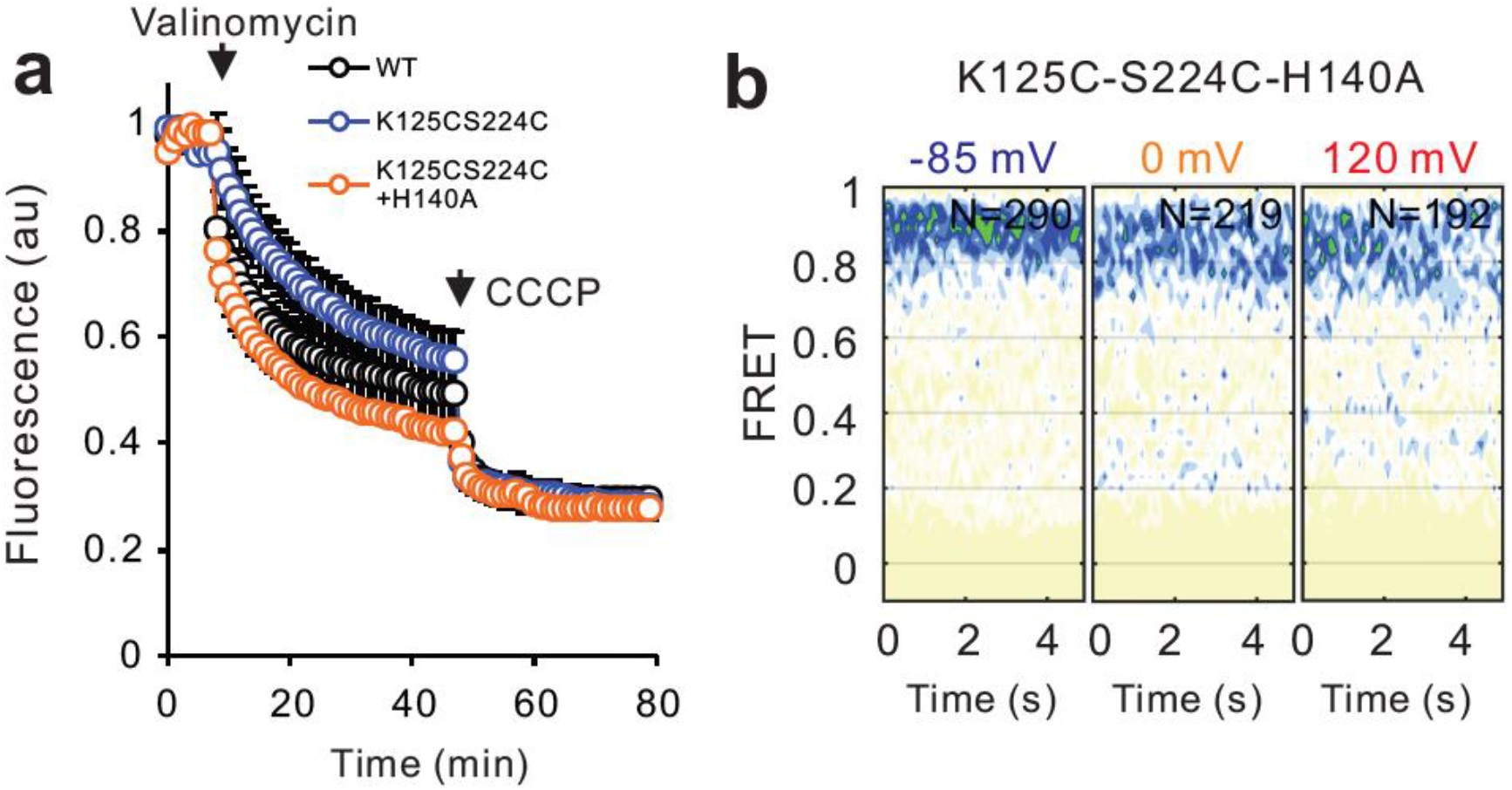
The activated conformations of the S4 segments in the hHv1 channel were captured by introducing the H140A mutation that abolishes pH inhibition. a. The liposome flux assays of the WT, K125CS224C and K125CS224C-H140A mutants, which are all functional to mediate proton uptake into liposomes. b. FRET histograms and contour maps from the K125CS224 labeling sites on the H140A mutation background.

